# NGSremix: A software tool for estimating pairwise relatedness between admixed individuals from next-generation sequencing data

**DOI:** 10.1101/2020.10.20.347500

**Authors:** Anne Krogh Nøhr, Kristian Hanghøj, Genis Garcia Erill, Ida Moltke, Anders Albrechtsen

## Abstract

Estimation of relatedness between pairs of individuals is important in many genetic research areas. When estimating relatedness, it is important to account for admixture if this is present. However, the methods that can account for admixture are all based on genotype data as input, which is a problem for low depth next-generation sequencing (NGS) data from which genotypes are called with high uncertainty. Here we present a software tool, NGSremix, for maximum likelihood estimation of relatedness between pairs of admixed individuals from low depth NGS data, which takes the uncertainty of the genotypes into account via genotype likelihoods. Using both simulated and real NGS data for admixed individuals with an average depth of 4x or below we show that our method works well and clearly outperforms all the commonly used state-of-the-art relatedness estimation methods PLINK, KING, relateAdmix, and ngsRelate that all perform quite poorly. Hence, NGSremix is a useful new tool for estimating relatedness in admixed populations from low-depth NGS data. NGSremix is implemented in C/C++ in a multi-threaded software and is freely available on Github https://github.com/KHanghoj/NGSremix.

## 1 Introduction

Estimation of genetic relatedness between pairs of individuals is important in many genetic research areas. For example, genetically related individuals are removed or accounted for in genome-wide association studies to avoid an inflated false positive rate. The relatedness between a pair of individuals is most often described using the concept of identity-by-descent (IBD), which is genomic identity due to recent common ancestry. To quantify the relatedness between a pair of individuals the summary statistics *R* = (*k*_0_*, k*_1_, *k*_2_) can be used, where *k*_0_, *k*_1_ and *k*_2_ are the proportion of the genome where a pair of individuals share 0, 1, or 2 alleles IBD, respectively. These statistics are useful because their expected values differ between different types of familial relationships, with the expected value of *k*_0_ being 1 for unrelated individuals and in general it is smaller the closer related two individuals are. For instance the expected values of *R* for sibling pair is (0.25, 0.5, 0.25) and for a parent offspring pair is (0, 1, 0). *R* can therefore be used to quantify how closely related two individuals are. There are also other IBD based summary statistics of relatedness, like the kinship coefficient, however we will here focus on *R*, because other such summary statistics can be calculated from it. For example, the kinship coefficient is simply 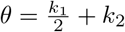.

Several methods exists for estimation of relatedness between a pair of individuals. When choosing a method to use it is important to consider what type of data is available, where the individuals are from and whether the individuals are admixed. Most current methods are based on the assumption that the individuals are from a homogeneous population [1, 2, 3, 4, 5], including the commonly used method implemented in PLINK [1]. If this assumption is violated the estimation of relatedness will be biased and relationships can be miss-classified [6, 7, 8]. To address this problem, several methods, such as RelateAdmix [6] and KING [9] have been developed. These methods are all developed to be applied to diallelic genotype data, like single nucleotide polymorphism (SNP) chip data. However, next-generation sequencing (NGS) data are becoming more common, and often this data is sequenced at low to medium depth, where genotype calling can come with significant biases [10]. This bias will be propagated into relatedness estimation and can lead to miss-classification of pairwise relatedness [11]. This issue can be avoided if instead of calling genotypes from low depth data one accounts for the genotype uncertainty by summing over all possible unobserved genotypes and weighting each of these using genotype likelihoods (GL). Several methods, like ngsRelate [11, 12], have used this approach to estimate pairwise relatedness from homogeneous populations from low depth sequencing data. However, none of these methods address relatedness estimates between individuals with admixed ancestry.

Here we present a maximum likelihood method, NGSremix, that can estimate relatedness (*R*) from low depth sequencing data when individuals have admixed ancestry. The method takes genotype likelihoods, admixture proportions and ancestral allele frequencies as input. The genotype likelihoods can be calculated using many software such as Samtools [13] and ANGSD [14], and ancestral allele frequencies can be estimated from clustering methods such a NGSadmix [15] or PCAngsd [16] assuming that the ancestral populations are discrete and there is NGS or genotype data available from a sufficient number of individuals. We also present a performance assessment of the method using both simulated data and sequencing data from the 1000 genomes project and compare its performance the commonly used state-of-the-art methods, PLINK [1], KING [9], relateAdmix [6], and ngsRelate [12] when applied to admixed individuals with low-depth NGS data. Importantly, the assessment shows that NGSremix works well and clearly outperforms all the other methods when there is admixture and you have low depth NGS data.

## 2 Method

### 2.1 The model

The main objective of the model is to enable maximum likelihood estimation of the relatedness coefficients (*R* = (*k*_0_*, k*_1_*, k*_2_)) for two individuals, 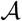 and 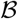, with ancestry from one or more of *K* different populations. We assume that we have NGS data from *M* variable diallelic sites for both individuals and denote this 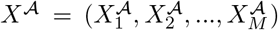 and 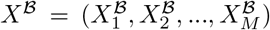. Furthermore, we assume that the paired ancestry proportions for both individuals, denoted 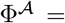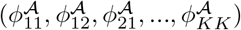 and 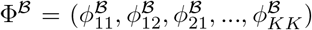 are known. Finally, we assume that for each site the ancestral allele frequencies for the *K* populations, *F*, are known. In practice we estimate the ancestral allele frequencies *F* using NGSadmix [15] and the paired ancestry proportion is estimated as described in Supplementary Material section 5.4. The paired ancestry proportions are simply the proportions of sites in the genome with a certain combination of ancestry, i.e. 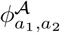 denotes the proportion of individual 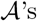 genome where the first allele is from population *a*_1_ and the second allele is from population *a*_2_.

Using this terminology we then write up the likelihood function for *R*. Since we assume 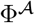, 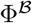 and *F* are known we have for simplicity not included them in this likelihood function. On the other hand, for each site the four alleles and their ancestry 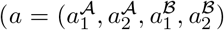, IBD status (*z* = (*z*_1_, *z*_2_)), and ordered genotypes 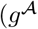 and 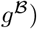 are not observed, see Figure 1. Therefore, we include these as latent variables and weight each possible value of these by their probability. Specifically, the likelihood function can be written as

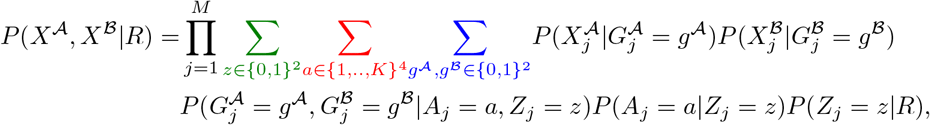

where *z* = (*z*_1_, *z*_2_) with *z*_1_ indicating whether allele 1 of individual 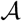 and 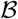 are IBD and *z*_2_ indicating whether the two individuals’ allele 2 are IBD. Furthermore, 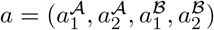 are the unobserved ancestral populations of the two individuals’ two alleles and 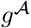 and 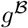 are the ordered genotypes for both individuals. Note, that we sum over all possible *ordered* genotypes such that both alleles from individual 1 can be IBD with any of the two alleles of individual 2, if they are from the same ancestry and have the same allelic state. 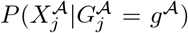 and 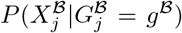 are GLs that represent the information from the sequencing data, which can be calculated as described in Section 2.2.3 e.g. using ANGSD [14]. The rest of the components of the likelihood function and a detailed description of how the likelihood function is derived are described in detail in Supplementary Material Section 5.2.

**Figure 1:**
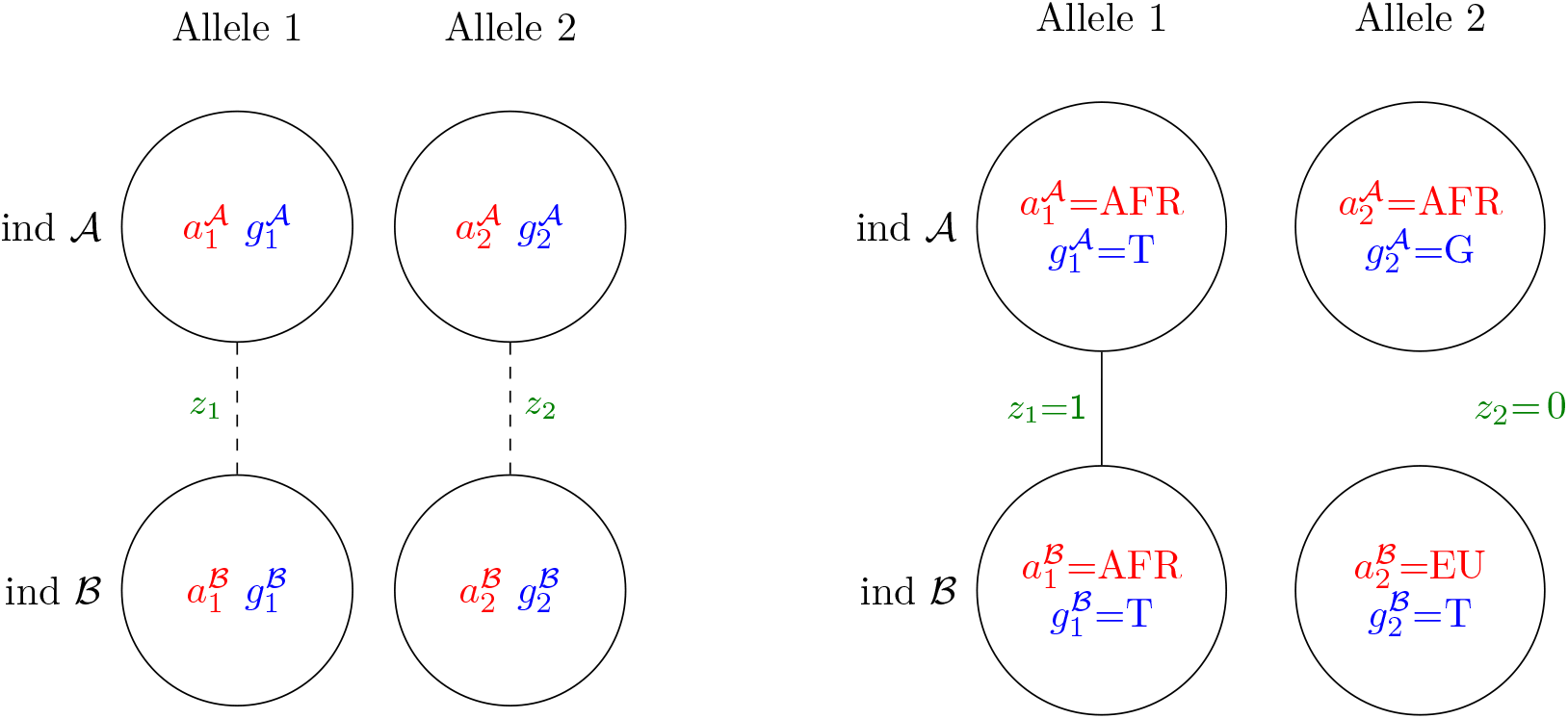
Left: Diagram of the unobserved state of the latent variables, *z*, *a*, 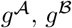, in a site. The circles represent the two alleles of each of the two individuals 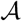 and 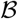. Specifically, for each of these individuals, e.g. individual 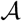, the first allele has an unobserved ancestry state 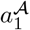, which can take any value among the *K* possible ancestral populations and an unobserved allelic state, 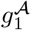. Similarly, the second allele has an unobserved ancestry state, 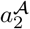, and an unobserved allelic state, 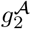, where 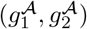 constitutes the unobserved ordered genotype, 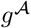, for the individual. Lines indicate the unobserved IBD states, i.e. whether the first alleles of the two individuals are IBD, *z*_1_, and whether the second alleles of the two individuals are IBD, *z*_2_. Right: Example diagram with a set of realized values for all the latent variables for two African American individuals with genotypes T/G and 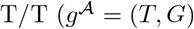 and 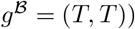, who share their first allele IBD (*z*_1_ = 1), which is possible because these alleles have the same African ancestry 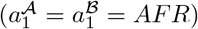 and the same allelic state 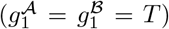. The second allele of the two individuals cannot be IBD (*z*_2_) because the two alleles originate from different ancestral populations 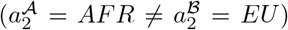 and/or because they have different allelic states 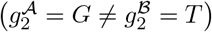. Note that we here used T and G instead of 0 and 1 as possible allelic states and AFR and EU instead of 1 and 2 as possible ancestral states to make the example more concrete.

To obtain maximum likelihood estimates of the paired ancestry Φ and the relatedness coefficients *R*, we are using an Expectation-Maximization (EM) algorithm. The EM algorithm is described in the Supplementary Material Section 5.3 and 5.4. For faster convergence, we implemented an accelerated EM algorithm using the squared iterative approach (S3) developed by Varadhan et al. [17]. The method is implemented in C/C++ in a multi-threaded software and is available on Github https://github.com/KHanghoj/NGSremix.

### 2.2 Simulation of data

Data were simulated to validate NGSremix and compare its performance with the existing methods: PLINK [1], KING [9], relateAdmix [6], and ngsRelate [12]. Using allele frequencies from Northern Europeans from Utah (CEU) and Yoruba in Ibadan, Nigeria (YRI) samples from the 1000 genomes project [18] we simulated NGS data with 100,000 diallelic sites for 10 pairs of individuals for each of 6 different relationship types: unrelated individuals (*R* = (1, 0, 0)), full siblings (*R* = (0.25, 0.5, 0.25), parent-offspring (*R* = (0, 1, 0)), half siblings (*R* = (0.5, 0.5, 0)), first cousins (*R* = (0.75, 0.25, 0)). The performance of the methods were evaluated using NGS data simulated based on 5 different average depths: 1x, 2x, 4x, 8x, and 16x. Data were generated by first simulating genotypes for each pair of individuals (see Section 2.2.1). Next, the genotypes were used to simulate NGS data (see Section 2.2.2) and finally GLs were calculated (section 2.2.3). Finally, we estimated ancestral allele frequencies from these GLs using NGSadmix [15], that accounts for genotype uncertainty, allowing for two ancestral sources and used these estimates along with the GLs as input to our method.

As input to ngsRelate we only used the GLs. For PLINK and KING that both require genotype data, we called genotypes from the calculated GLs by choosing the genotype with the highest GL and used these as input. For RelateAdmix that requires genotypes as well as admixture proportions and ancestral allele frequencies, we first used the called genotypes to estimate admixture proportions and ancestral allele frequencies using using ADMIXTURE [19] and then used these estimates as well as the called genotypes as input.

#### 2.2.1 Simulation of genotypes

For unrelated pairs we simulated genotypes without linkage disequilibrium by first randomly sampling an admixture proportion between the two ancestral populations (CEU and YRI) of 0, 0.25, 0.5, 0.75 or 1 for each individual. Then for each site the ancestry of the alleles for each of the two individuals were sampled using the admixture proportions. Finally, the genotypes were sampled based on the sampled ancestral populations at that site and the allele frequencies from the 1000 genomes project [18]. For the remaining relationship types, haplotypes were simulated by first simulating genotypes for an appropriate number of unrelated admixed founder individuals using the same approach as for the unrelated pairs. Next, we simulated data for the relevant pair by simulating offspring of these founders (and in some cases their offspring) using the relevant pedigree. E.g. for siblings we simulated data for two unrelated founder individuals (parents) and then simulated the siblings by simulating two offspring from these parents. In all cases, genotypes for an offspring of two parents were simulated at each site by randomly sampling one allele from each parent.

#### 2.2.2 Simulation of NGS data

Based on the simulated genotypes, NGS data was generated for each individual as follows. First, the sequencing depth *d* at each site was sampled from a Poisson distribution with mean equal to the specified average sequencing depth. Next, *d* bases were sampled at each site j, *X_j_* = (*b*_1_*, b*_2_*,…, b_d_*), based on the individual’s genotype and a per base sequencing error probability, *ϵ*, of 0.005. In case of a sequencing error, the base is replaced with the other possible base at the site.

#### 2.2.3 Calculation of genotype likelihoods

GLs were calculated using the model described by Mckenna et al. [20]. However, for simplicity we assume that there is only two alleles. At each site *j* and for each of the genotypes, 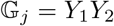 where *Y*_1_, *Y*_2_ ∈ {0, 1}, we calculated the likelihood of observing the NGS data *X_j_* = (*b*_1_, *b*_2_,…, *b_d_*), as

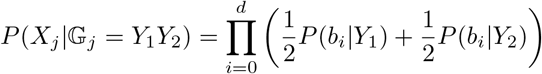

where

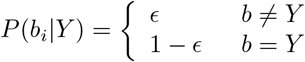

Note, that this model was originally described for unordered genotypes, but that the genotype likelihoods for ordered genotypes are the same as for unordered genotypes.

### 2.3 Real data

We also tested NGSremix on real data from 268 individuals from the 1000 genomes project [18]. These individuals have all been sequenced at low depth, however, they have also been genotyped using SNP chips and many of them have been sequencing with high depth exome and/or whole genome sequencing. Therefore, we also have a high quality genotype data set to compare the low depth sequencing data to. Specifically, we analysed individuals from the admixed population of Americans of African Ancestry (ASW, n=61). We also included some European individuals (CEU, n=99) and some African individuals (YRI, n=108) in order to estimate accurate admixture proportions and ancestral allele frequencies for the admixed ASW individuals. To assess performance on low depth NGS data, we analysed the low depth NGS data for these samples which have a median depth around 4x depth of coverage. First, we calculated genotype likelihoods (GL) (-gl 2-Q 30 -q 30) for all individuals using ANGSD [14]. When doing so we restricted the analysis to polymorphic sites in the high quality genotype data for these three populations with a minor allele frequency of 0.05, followed by LD pruning (PLINK1.90 –indep-pairwise 50 10 0.1) [21]. A total of 329.453 sites were used for downstream analysis. Next, we estimated the ancestral allele frequencies using NGSadmix [15], that accounts for genotype uncertainty, allowing for two ancestral sources. Finally, we applied NGSremix to the GL data, ancestry proportions, and ancestral allele frequencies to estimate the relatedness coefficients (*R*) for all pairs of individuals.

For comparisons using PLINK, KING and relateAdmix, we called genotypes from the low depth NGS data by choosing the maximal genotype likelihood for each site and each individual and used these as input. Furthermore, for relateAdmix we estimated ancestry proportions and ancestral allele frequencies from the called genotypes using ADMIXTURE [19] and used these as additional input.

To validate the results obtained from the low depth sequencing data, we performed a set of secondary analyses using the high quality genotype data for the same individuals using the same set of genomic sites.

## 3 Results

### 3.1 Performance assessment using simulated data

We first assessed the performance of NGSremix on simulated data. We simulated genotype data for 6 different relationship types and various admixture scenarios based on allele frequencies from the CEU and YRI 1000 Genomes population data. From these genotype datasets we then simulated NGS data with depths 1x, 2x, 4x, 8x, and 16x. Finally, we estimated *k*_0_, *k*_1_, and *k*_2_ values for the simulated NGS data for all pairs of individuals (N=7140) using our NGSremix as well as the four commonly used state-of-the-art methods ngsRelate, relateAdmix, PLINK, and KING. Since KING estimates kinship and not *R*, the kinship coefficient for NGSremix and KING were also compared. When analysing the simulated data with average depths of 8x or 16x, all methods can distinguish between full siblings, half-siblings, and parent-offspring (Supplementary Material Figures S1 and S2). However, only our NGSremix and relateAdmix obtain accurate estimates and allows clear separation of unrelated individuals from first-cousin. For simulated NGS data with an average depth of 4x, i.e. low depth NGS data, NGSremix is the only method that gives accurate results and allows the different relationship types to be distinguished from each other (Figure 2). ngsRelate give somewhat reasonable estimates for the first degree relationships, however cannot separate the unrelated, second-cousin, and first-cousin. relateAdmix on the other hand, can for the most part distinguish between the relationships, but the relatedness coefficient estimates cannot be interpreted as IBD fractions making it difficult to use them e.g. for relationship classification. The last two methods, PLINK and KING, performs worse in this scenario with estimated relatedness and kinship coefficients that are difficult to interpret. Similar results are obtained at lower depths (Supplementary Material Figures S3 and S4); even on 1x data our method performs fairly well, especially when increasing the number of sites, and is able to differentiate all relationship types except from unrelated and second cousins, see Supplementary Material Figure S5.

**Figure 2:**
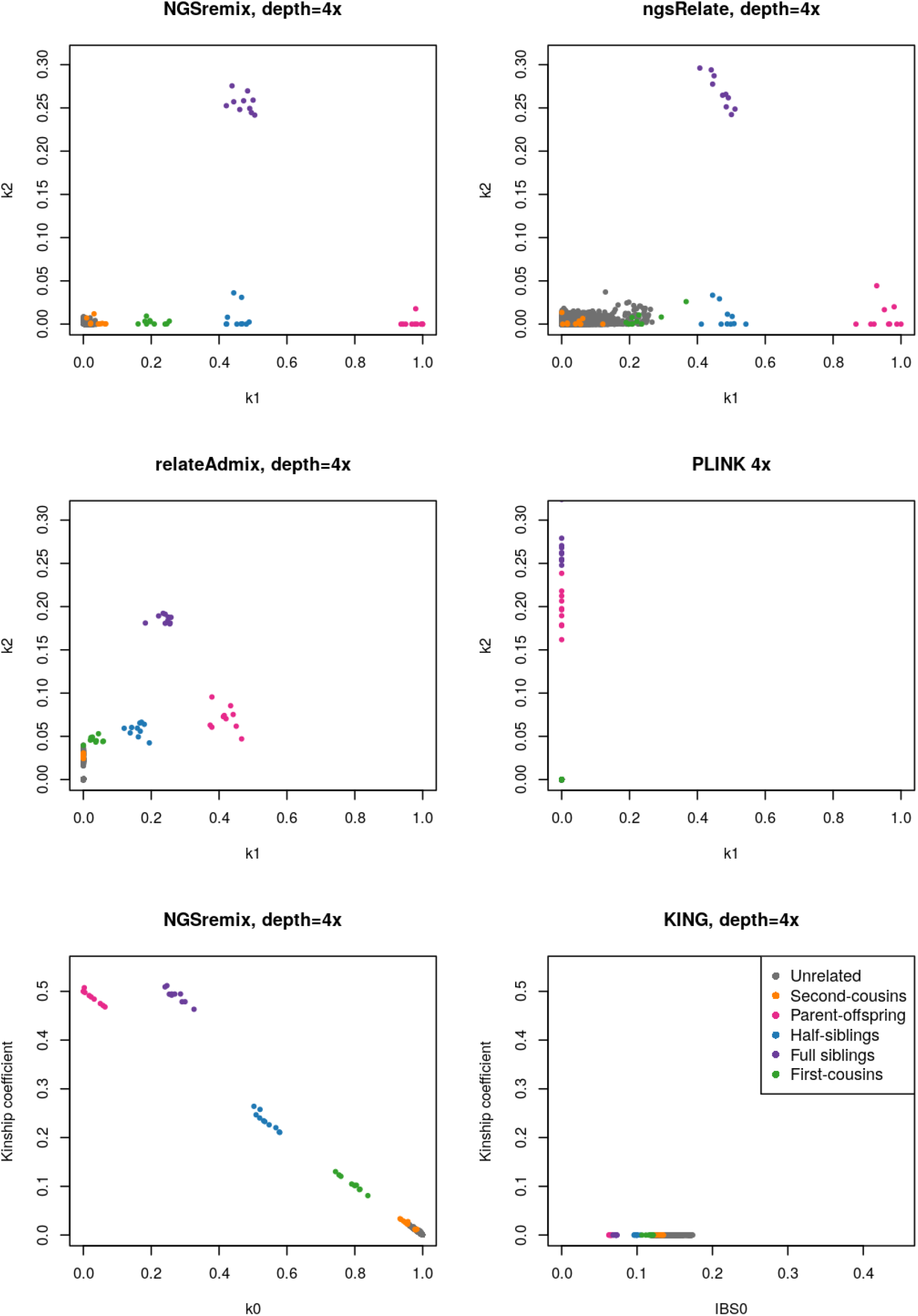
Estimated *k*_1_ and *k*_2_ or kinship for 140 simulated admixed individuals with 6 relationship types (10 of each type of related pairs) and an average depth of 4x. Plots of estimated R values are visualized for NGSremix, ngsRelate, reateAdmix, and PLINK. Plots of the kinship coefficient for NGSremix and KING are provided since KING only estimates kinship

### 3.2 Performance assessment using real data

To further assess the performance of NGSremix we also applied the same methods to data from the admixed African American individuals from the 1000 genomes project. First, we applied the methods to high quality genotype data for these individuals to obtain estimates that are as close to the truth as possible, see Supplementary Material Figure S6. NGSremix and relateAdmix that are both designed to take admixture into account, gave close to the same results and identified 5 parent-offspring pairs, 1 pair of half siblings and 2 pairs of individuals as first-cousins. These results are consistent with previous reports for these pairs [22]. In contrast PLINK, ngsRelate and KING showed larger variance over estimates *k*_0_ and *k*_1_ and while they identify the same 5 parent-offspring pairs and the half-sibling pair, the results for the 2 first cousin pairs are either difficult to interpret (due to high *k*_2_ estimates) or indicate a first cousin relationship and/or can not be distinguished from the unrelated pairs.

We next applied the methods to low-depth sequencing data (around 4x) from the same individuals. The results showed that our method is the only method that is able to identify the same related individuals as identified from the high quality genotype data by both NGSremix and relateAdmix (Figure 3. ngsRelate have similar, suboptimal results as it did when it was applied to the high quality genotype data, while relateAdmix, PLINK and KING performed markedly worse. Hence, our method clearly outperforms the existing methods when applied on low-depth NGS data, not only for simulated data but also for real data.

**Figure 3:**
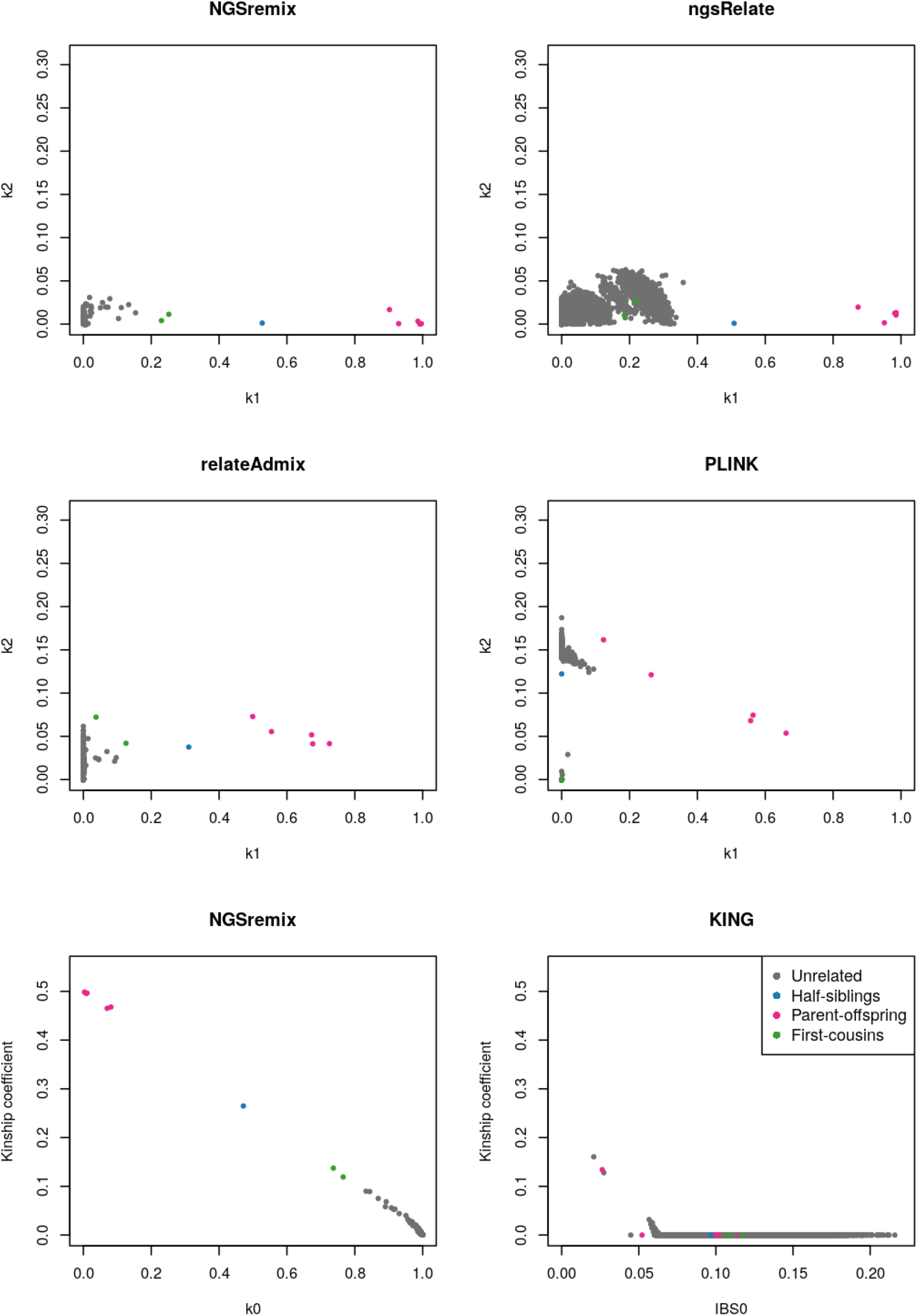
Inferred relatedness, *k*_1_ and *k*_2_ or kinship, for 168 admixed African Americans from the 1000 genomes project sequenced at low depth. Plots of estimated R values are visualized for NGSremix, ngsRelate, relateAdmix, and PLINK. Plots of the kinship coefficient for NGSremix and KING are provided since KING estimates kinship and not *R*.

## 4 Discussion

We have presented a new maximum likelihood based software tool, NGSremix, for estimation of pairwise relatedness between admixed individuals sequenced at low depth. Using simulated data and data from the 1000 genomes project, we have showed that our method is superior when used on admixed individuals sequenced at low depth compared to current state-of-the-art methods: PLINK, KING, relateAdmix, and ngsRelate. NGSremix’s accurate estimates of relatedness are due to the fact that it takes GLs as input instead of called genotypes, which is an advantage for low-depth NGS data, where genotypes cannot be called with high certainty. Secondly, NGSremix handles admixture by estimating paired ancestry proportions and including these in the model.

From the results of the simulations, we saw that the state-of-the-art methods that rely on genotypes as input could separate full-siblings, parent-offspring, and half-siblings when the average depth was 16x. However, KING, ngsRelate and PLINK gave highly inaccurate relatedness/kinship estimates for the sibling and unrelated pairs. In contrast, relateAdmix [6] provided accurate estimates close to similar to the results of NGSremix, which is not a surprise as relateAdmix account for admixture and is methodically very similar to NGSremix. This shows that calling genotypes is a good option if the depth is high enough to call genotypes correctly and when an appropriate method that takes admixture into account is used. However, when lowering the simulated depth to 8x the genotype-based methods, including relateAdmix, obtained less accurate results compared to those based on 16x data. For depths below 8x, the uncertainty of genotype calls reached a level where the genotype-based methods began to give nonsensical results e.g. for 4x data did PLINK [21] and KING [9] estimate all *k*_1_ coefficients and kinship coefficients to zero for all pairs of individuals regardless of relationship type. Thus, our results show that calling genotypes for low depth NGS data can greatly bias estimation of pairwise relatedness. This is not the case when using genotype likelihoods, because they contain all information about the uncertainty of the genotype, while this information is lost when calling a genotype. Even for simulated data of depth 1x our method provided sensible results, however, albeit more noise was observed. We can not make an exact statement about how low a depth our method can handle. Since it will depend on number of SNPs, how admixed the individuals are, and the number of individuals. However, we expect it will not work well with lower depth since observing reads containing each individuals two alleles would be required at least at some sites.

All the tested methods in this paper, including NGSremix, has a series of limitations. First, the estimated relatedness from the real data can deviate from the expectation for all relationship pairs except for parent-offspring and monozygotic twins, e.g. the kinship coefficients for a pair of full-siblings is expected to be 0.5, but will often deviate from this exact value. This is due to biological variance during the recombination process, where a pair can share more/less IBD segments than expected [23] with variable variance depending on the relationship [24]. This can in some cases make it challenging to disentangle the actual degree of a pair of relatives. Second, all the methods either assume allele frequencies are from discrete homogeneous populations or in the case of KING assumes that each pair of individuals is from the same homogeneous populations. Third, for all of the methods except for KING the number of individuals included in the analysis is important since they are based on the assumption that allele frequencies can obtained. For NGSremix and relateAdmix the number of individuals is even more important because these methods are based on the assumption that the ancestral allele frequencies and admixture proportions can be accurately estimated using clustering methods such a NGSadmix [15] or PCAngsd [16] can be used. The exact number of individuals per population needed in order to obtain accurate estimates from these method will depend on the number of sites, the admixture patterns and the population degree of populating differentiation. However, having less than 5 individuals representing a population will lead to inaccurate estimated [25]. Thus NGSremix cannot be applied to studies were only a few individuals have been sequenced. Instead it requires sequencing or genotype data for multiple individuals representing each ancestral population. Lastly, NGSremix assumes that there is no inbreeding and that the markers are independent i.e. no linkage disequilibrium. These assumptions are also shared with the other methods with the exception of ngsRelate which allows for arbitrary inbreeding patterns [12]. With regards to the assumption of no linkage disequilibrium then all of the methods can still be applied with data which has not been pruned. In this case, the models should be viewed as composite likelihood models which will have a different interpretation of the likelihood value. However, it will not affect the expected maximum likelihood point estimate [26].

In conclusion we have presented a maximum likelihood method for estimating relatedness for low depth sequencing data that can be applied to admixed individuals. In simulations and real data from low depth sequencing of admixed individuals the method outperforms other methods and gives reasonable results down to 1x sequencing.

## 5 Supplementary Material

### 5.1 Notation

Throughout the Supplementary Material we will use the following notation:

*M* Number of diallelic sites
*K* Number of ancestral populations
*R* Relatedness coefficient with *R* = (*k*_0_, *k*_1_, *k*_2_) where *k_i_* is the proportion of the genome where individuals 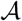 and 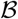 share *i* alleles IBD
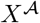 NGS data from individual 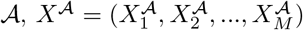
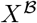 NGS data from individual 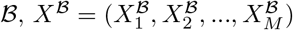
*F* Matrix of ancestral allele frequencies for *K* populations for *M* sites *F* = (*F*_1_, *F*,…, *F_M_*) with *F_j_* = (*F_j_*_1_, *F_j_*_2_,…, *F_jK_*)
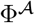 Paired ancestry proportions for individual 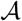, with paired ancestry from *K* different populations. where 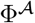 contains *K*^2^ entries, 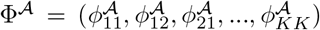, where 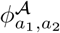 denotes the proportion of the genome where the first allele is from population *a*_1_ and the second allele is from population *a*_2_.
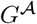 Latent variable of the ordered genotypes from individual 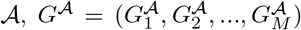 each with realization 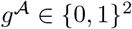
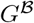 Latent variable of the ordered genotypes from individual 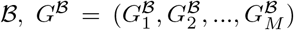 each with realization 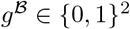
*Z* Latent variable of the IBD state of allele 1 and 2, *Z* = (*Z*_1_, *Z*_2_,…, *Z_M_*) each with realization *z* = (*z*_1_, *z*_2_) ∈ {0, 1}^2^
*A* Latent variable of the ancestral state of the two individuals’ pair of alleles *A* = (*A*_1_, *A*_2_*,…, A_M_*) each with realization 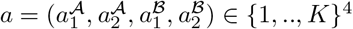
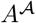 Latent variable of the ancestral state of individual 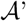 pair of alleles 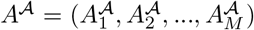 each with realization 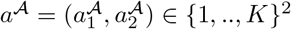

### 5.2 Derivation of the likelihood function

We want to estimate relatedness coefficients, *R* = (*k*_0_, *k*_1_, *k*_2_), for two individuals, 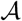 and 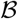, with ancestry from *K* different populations using a maximum likelihood approach. *R* = (*k*_0_, *k*_1_, *k*_2_) denote the fractions of the genome where the pair of individuals share 0, 1, and 2 alleles IBD (Σ*k_i_* = 1) and in the following we derive the likelihood function that we use to estimate these.

For each individual we assume we have NGS data from *M* variable diallelic sites, 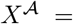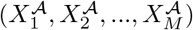 and 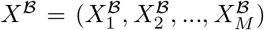. We let 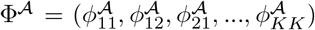 and 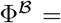 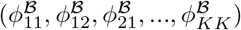 denote the paired ancestry proportions of the two individuals and *F* denote the allele frequencies of the *K* ancestral populations for each of the *M* sites. We assume that both Φ and *F* are known and for simplicity we have not included them in the likelihood function. For each site, the genotypes of the two individuals are unobserved. Therefore, we include ordered genotypes for both individuals, 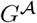 and 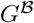, as latent variables in the model weighted by their probability. The likelihood function based on NGS data can thus be written

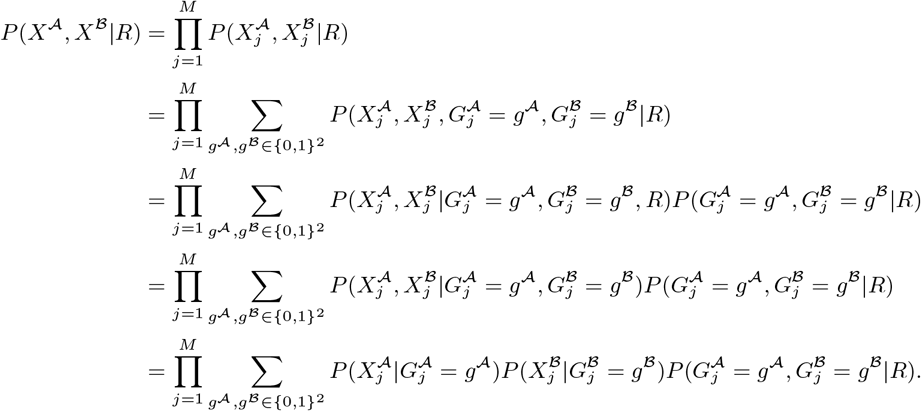

Here we assumed that the sites are independent and that the probability of observing the NGS data is independent between the two individuals conditioned on the genotype being known. We end up with a product of the genotype likelihoods of the two individuals and the probability of the genotypes given the relatedness coefficients. The latter is the same likelihood that is used by relateAdmix [6].

Building on this model, we can introduce the ancestry and the IBD state in the model. For each site, the four alleles’ IBD status and ancestry are also not observed. Therefore, we also include these as latent variables in the model weighted by their probability. Specifically, we let *z* = (*z*_1_, *z*_2_) denote IBD status, where *z*_1_ indicates whether allele 1 of individuals 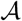 and 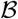 are IBD and *z*_2_ indicates whether allele 2 in individuals 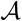 and 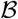 are IBD. Also, we let 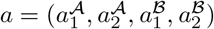 denote the unobserved ancestral populations of the two individuals’ alleles. When including *z* and *a* the likelihood becomes

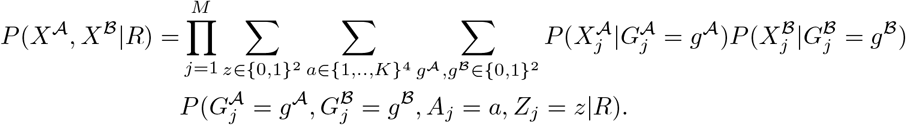

The last term can be be rewritten to

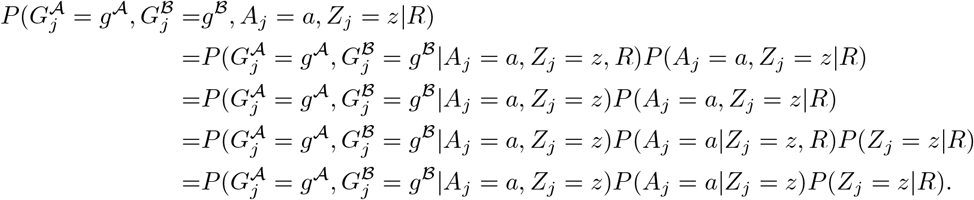

After rewriting the last term, the likelihood will finally become:

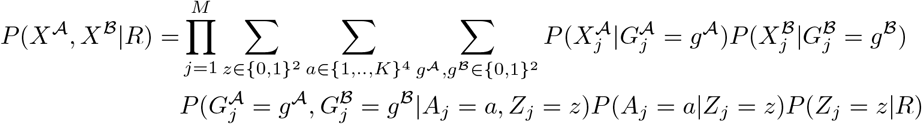

where 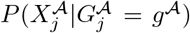 and 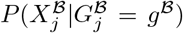 are GLs and they can be calculated as described in Section 2.2.3. The factor 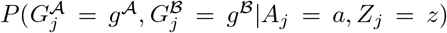 is described in Table S1, *P* (*A_j_* = *a*|*Z_j_* = *z*) is described in Table S2, and *P* (*Z_j_* = *z*|*R*) are given by:

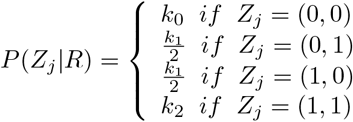

### 5.3 EM algorithm

The relatedness coefficients *R* = (*k*_0_, *k*_1_, *k*_2_) are estimated by finding the maximum likelihood (ML) using an EM algorithm. The expectation of the relatedness for each site *j* is:

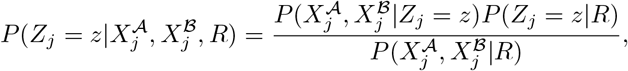

where

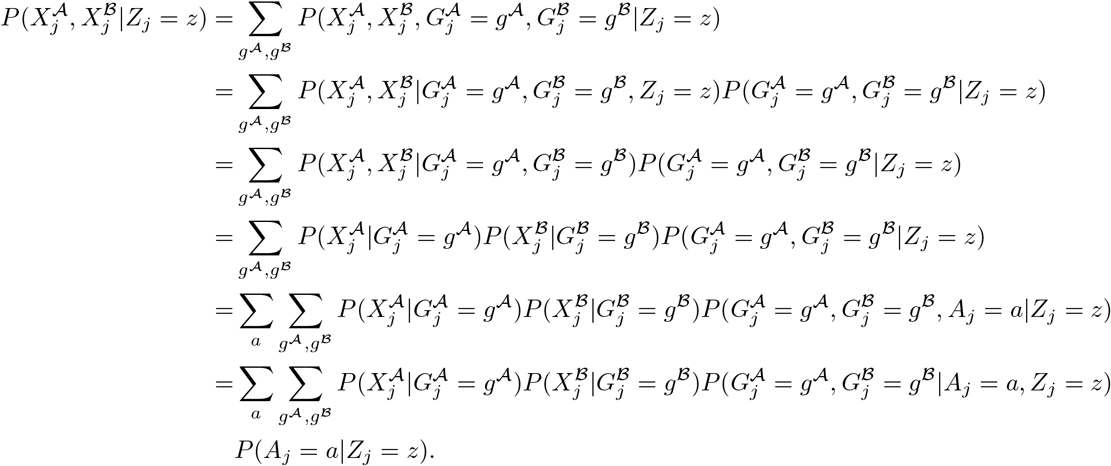

The derived likelihoods can be found in Table S1 and S2. The initial starting parameters, 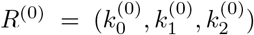, of the EM algorithm are random. For each *i*th iteration of the EM algorithm the parameters are updated to *R*^(*i*+1)^, using the current parameters (*R*^(*i*)^):

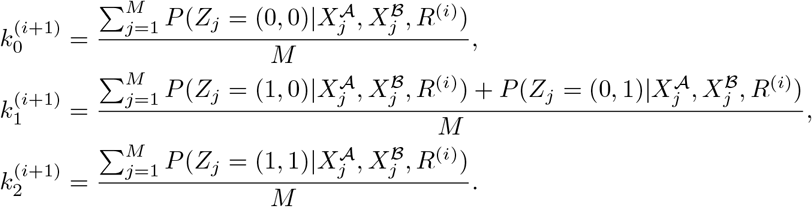

The EM algorithm will continue to update the relatedness coefficients until the algorithm converges. The algorithm converges when the root sum squared difference between *R*^(*i*)^ and *R*^(*i*+1)^ is less than 1*e^−^*^6^.

### 5.4 Estimation of paired ancestries

A recent admixture event, such as an offspring of parents from different population, will cause a dependency of the ancestry of the offspring’s two alleles. At each site in the genome the offspring will have exactly one allele from each of the two populations. In order to accommodate this assumption we estimate paired ancestry for each individual. For individual 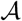 with ancestry from *K* populations, we define a vector 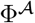 containing *K*^2^ entries, 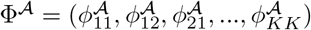, where 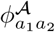 gives the proportion of sites from the individual’s genome where the first allele has ancestry from ancestral population *a*_1_ and the second one from *a*_2_. Although the pairs of ancestries are ordered we assume that they are symmetric i.e. 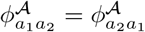. We assume that we know, or that we have previously estimated, the ancestral allele frequencies in each population, *F*. The likelihood of observing the sequencing data 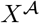 for a single individual, *M* sites and *K* ancestral populations is then

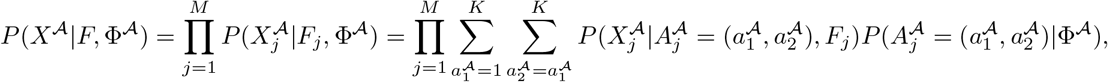

where the probability of a paired ancestry is directly specified by 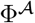,

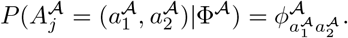

The probability of the NGS data 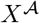 at site *j* given that site has paired ancestry 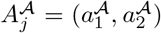 is

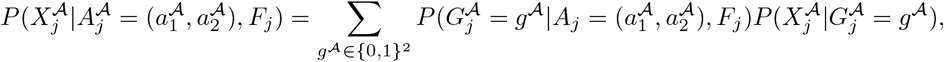

where

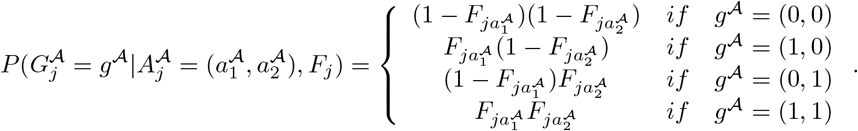

#### 5.4.1 EM for Φ

In the following 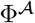 and 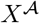 refer respectively to the paired ancestry proportions and the sequencing data for a single individual 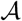. We use an expectation maximization (EM) algorithm to obtain a maximum likelihood estimate 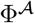. We denote the *i*th iteration of the estimated pairwise ancestry proportion Φ^(*i*)^. Based on this estimate, the expectation of the total number of sites with paired ancestries 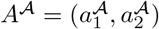 is given by

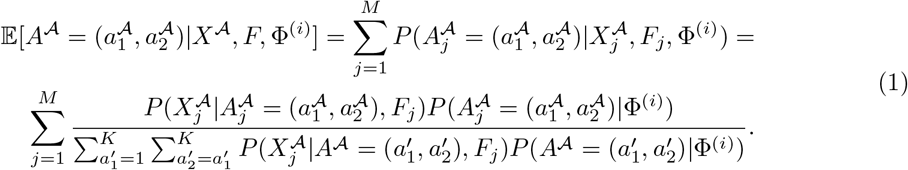

Then we can update the estimate of Φ by calculating each of its entries as

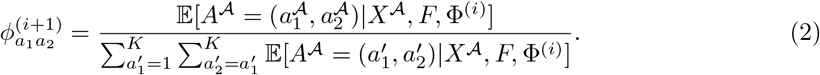

Finally, we calculate the new expectation given Φ^(*i*+1)^ with equation (1) and use the expectation to calculate a new estimate with equation (2) iterating until convergence. A proof that this is an EM for a similar case can by found in [15].

### 5.5 Supplementary figures

**Figure S1:**
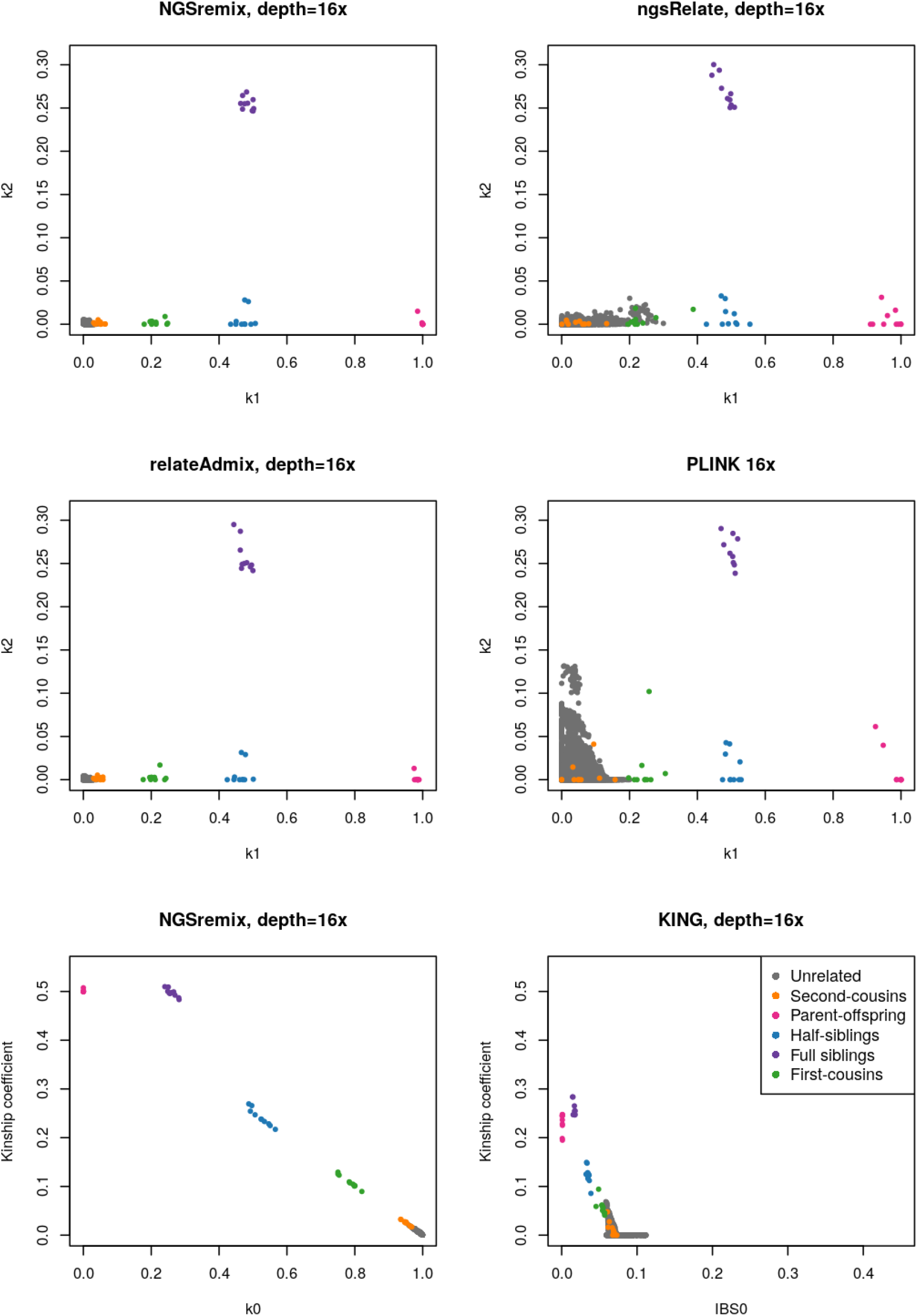
Estimated *k*_1_ and *k*_2_ or kinship for 140 simulated admixed individuals with 6 relationship types (10 of each type of related pairs) and an average depth of 16x. Plots of estimated R values are visualized for NGSremix, ngsRelate, reateAdmix, and PLINK. Plots of the kinship coefficient for NGSremix and KING are provided since KING only estimates kinship

**Figure S2:**
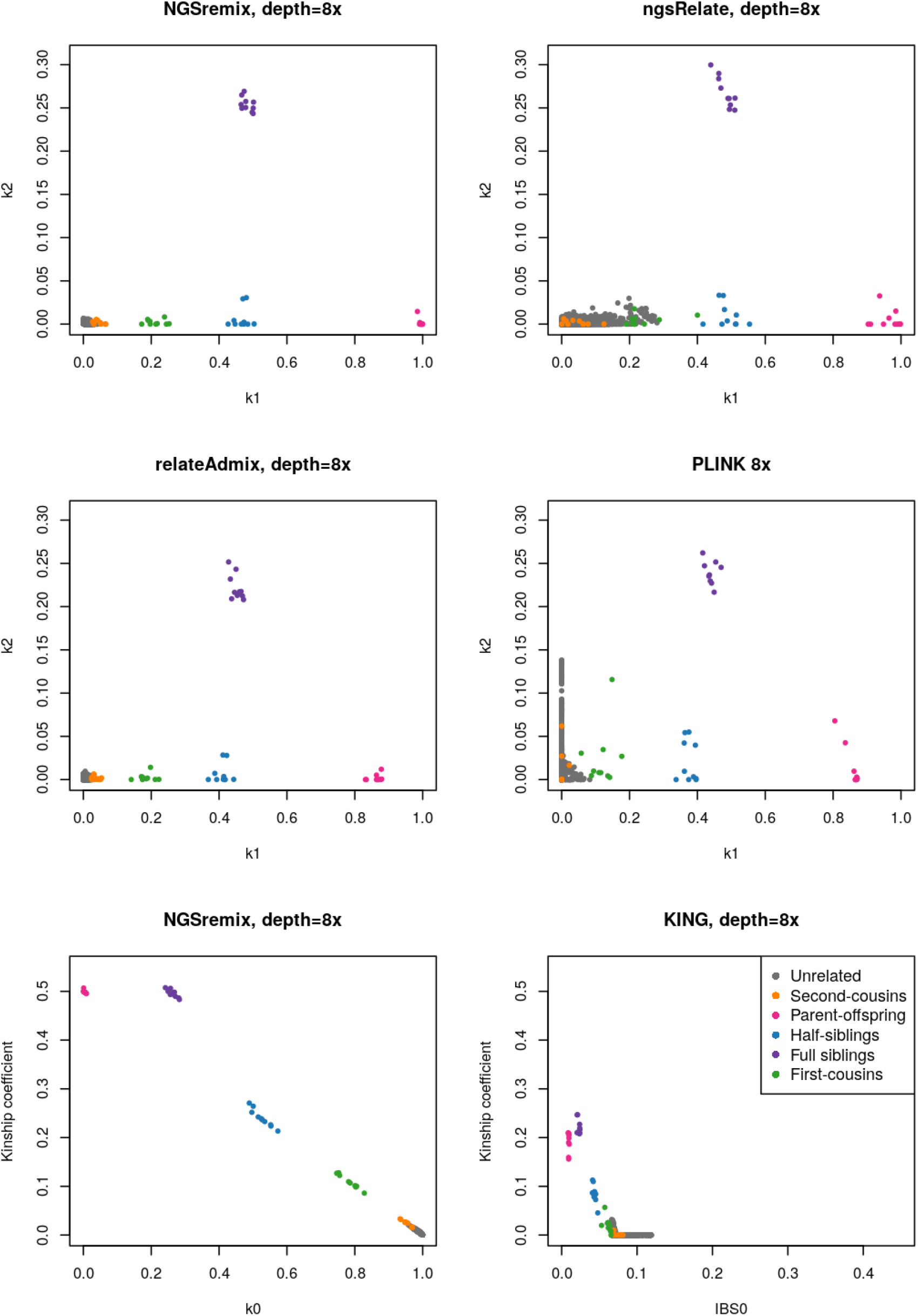
Estimated *k*_1_ and *k*_2_ or kinship for 140 simulated admixed individuals with 6 relationship types (10 of each type of related pairs) and an average depth of 8x. Plots of estimated R values are visualized for NGSremix, ngsRelate, reateAdmix, and PLINK. Plots of the kinship coefficient for NGSremix and KING are provided since KING only estimates kinship

**Figure S3:**
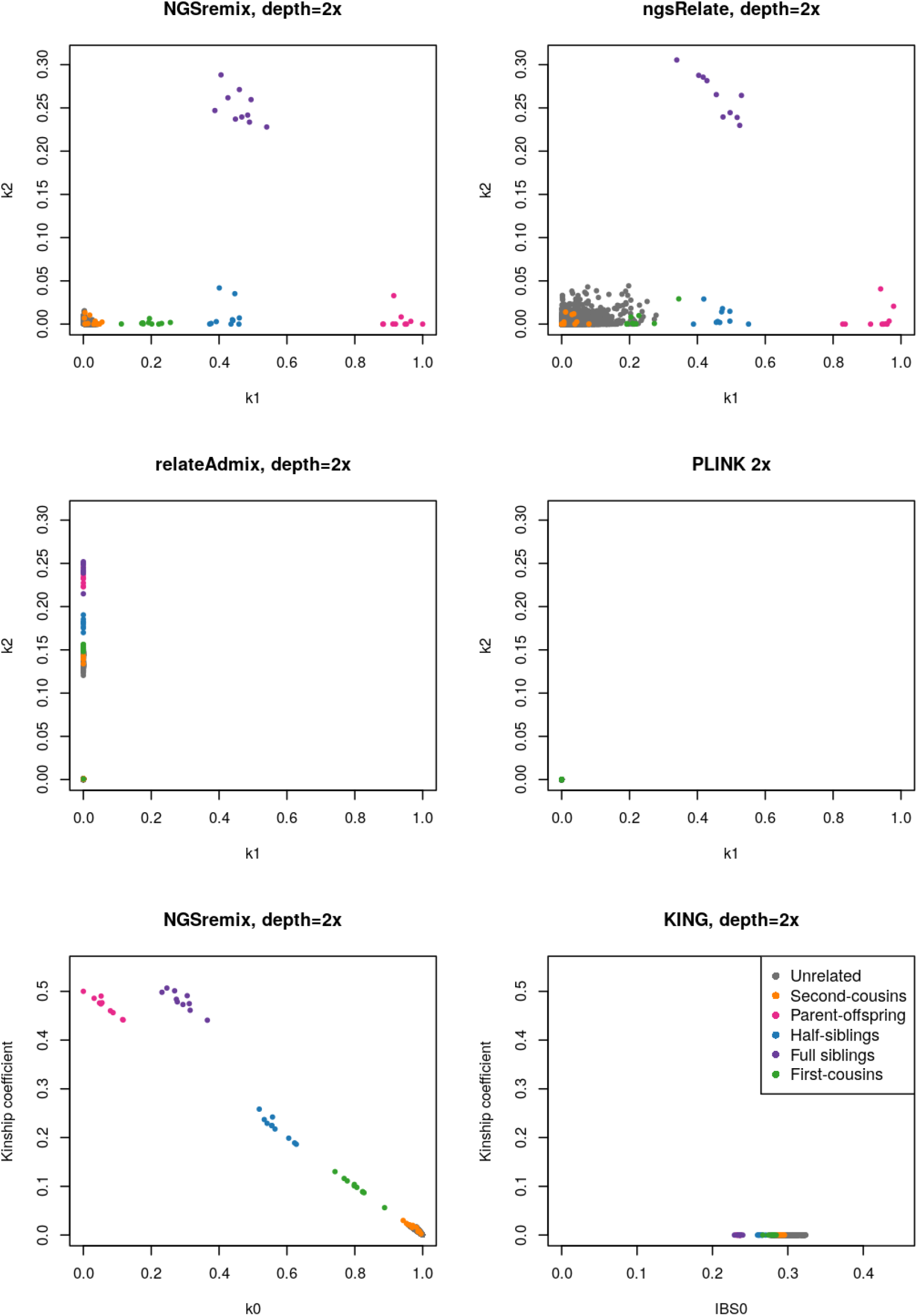
Estimated *k*_1_ and *k*_2_ or kinship for 140 simulated admixed individuals with 6 relationship types (10 of each type of related pairs) and an average depth of 2x. Plots of estimated R values are visualized for NGSremix, ngsRelate, reateAdmix, and PLINK. Plots of the kinship coefficient for NGSremix and KING are provided since KING only estimates kinship

**Figure S4:**
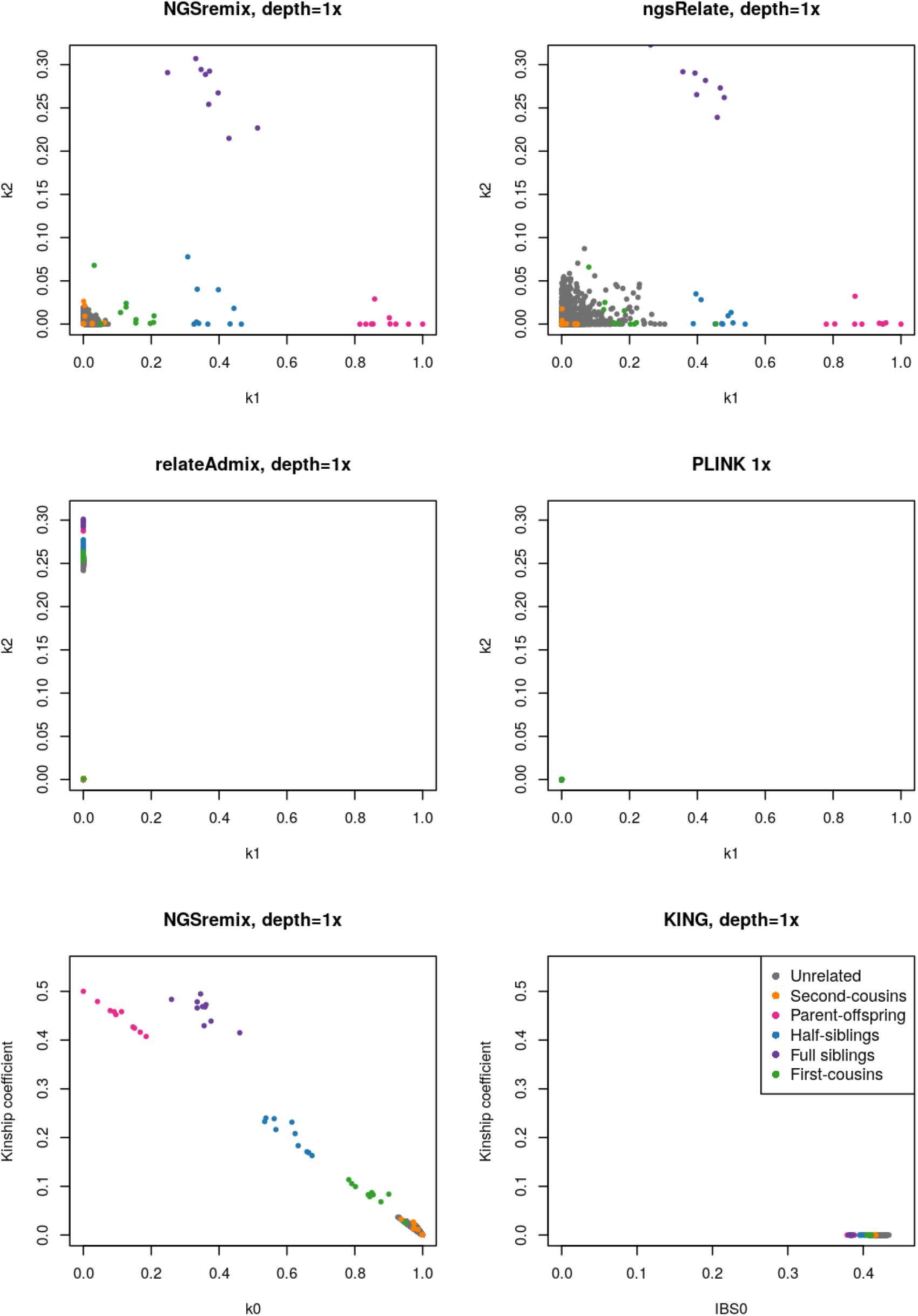
Estimated *k*_1_ and *k*_2_ or kinship for 140 simulated admixed individuals with 6 relationship types (10 of each type of related pairs) and an average depth of 1x. Plots of estimated R values are visualized for NGSremix, ngsRelate, reateAdmix, and PLINK. Plots of the kinship coefficient for NGSremix and KING are provided since KING only estimates kinship

**Figure S5:**
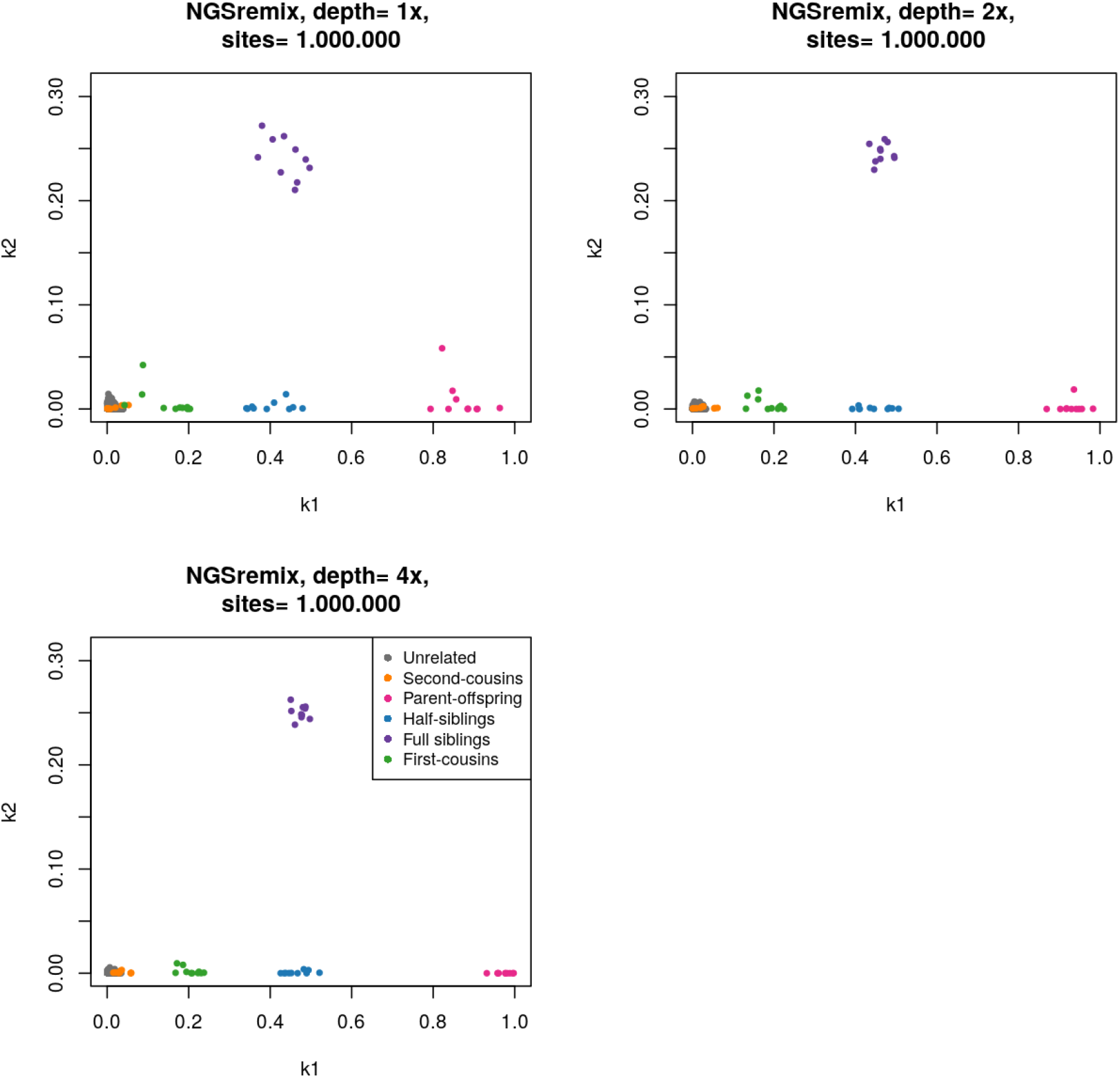
*k*_1_ and *k*_2_ values estimated using NGSremix for 140 simulated admixed individuals with 1.000.000 sites and 6 relationship types (10 of each type of related pairs) and an average depth of 1x, 2x, or 4x.

**Figure S6:**
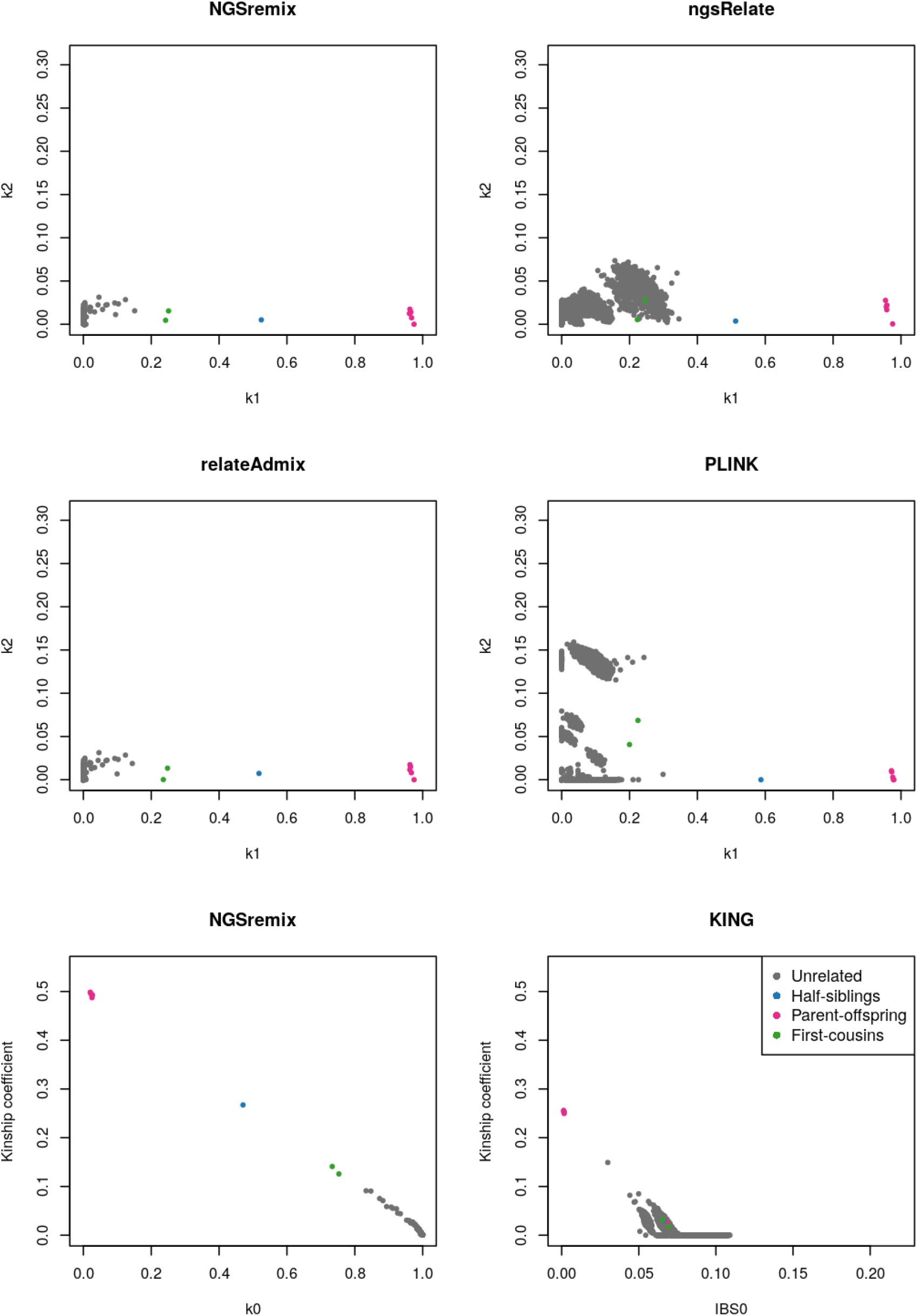
Inferred relatedness, *k*_1_ and *k*_2_ or kinship, for 168 admixed African Americans from the 1000 genomes project sequenced at high depth. Plots of estimated R values are visualized for NGSremix, ngsRelate, reateAdmix, and PLINK. Plots of the kinship coefficient for NGSremix and KING are provided since KING estimates kinship and not R.

### 5.6 Supplementary tables

**Table S1:** Calculation of 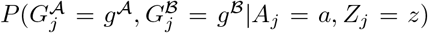 for all possible combinations of 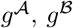, *z*_1_, *z*_2_, and 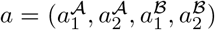. From the ancestral allele frequency matrix *F* we extract the allele frequency from site *j* of allele 0 from population *k* and define as *p_k_* = *F_jk_*. The allele frequency of allele 1 in population *k* we define as *q_k_* = 1 − *p_k_*.

| $g^A g^B$ | $z_1 = 0 \ z_2 = 0$ | $z_1 = 0 \ z_2 = 1$ | $z_1 = 1 \ z_2 = 0$ | $z_1 = 1 \ z_2 = 1$ |
| --- | --- | --- | --- | --- |
| 00 00 | $p_{a_1^A} p_{a_2^A} p_{a_1^B} p_{a_2^B}$ | $p_{a_1^A} p_{a_2^A} p_{a_1^B}$ | $p_{a_1^A} p_{a_2^A} p_{a_2^B}$ | $p_{a_1^A} p_{a_2^A}$ |
| 00 01 | $p_{a_1^A} p_{a_2^A} p_{a_1^B} q_{a_2^B}$ | 0 | $p_{a_1^A} p_{a_2^A} q_{a_2^B}$ | 0 |
| 00 10 | $p_{a_1^A} p_{a_2^A} q_{a_1^B} p_{a_2^B}$ | $p_{a_1^A} p_{a_2^A} q_{a_1^B}$ | 0 | 0 |
| 00 11 | $p_{a_1^A} p_{a_2^A} q_{a_1^B} q_{a_2^B}$ | 0 | 0 | 0 |
| 01 10 | $p_{a_1^A} q_{a_2^A} q_{a_1^B} p_{a_2^B}$ | 0 | 0 | 0 |
| 01 00 | $p_{a_1^A} q_{a_2^A} p_{a_1^B} p_{a_2^B}$ | 0 | $p_{a_1^A} q_{a_2^A} p_{a_2^B}$ | 0 |
| 01 01 | $p_{a_1^A} q_{a_2^A} p_{a_1^B} q_{a_2^B}$ | $p_{a_1^A} q_{a_2^A} p_{a_1^B}$ | $p_{a_1^A} q_{a_2^A} q_{a_2^B}$ | $p_{a_1^A} q_{a_2^A}$ |
| 01 11 | $p_{a_1^A} q_{a_2^A} q_{a_1^B} q_{a_2^B}$ | $p_{a_1^A} q_{a_2^A} q_{a_1^B}$ | 0 | 0 |
| 10 00 | $q_{a_1^A} p_{a_2^A} p_{a_1^B} p_{a_2^B}$ | $q_{a_1^A} p_{a_2^A} q_{a_1^B}$ | 0 | 0 |
| 10 01 | $p_{a_1^A} q_{a_2^A} q_{a_1^B} p_{a_2^B}$ | 0 | 0 | 0 |
| 10 11 | $q_{a_1^A} p_{a_2^A} q_{a_1^B} q_{a_2^B}$ | 0 | $q_{a_1^A} p_{a_2^A} q_{a_2^B}$ | 0 |
| 10 10 | $q_{a_1^A} p_{a_2^A} q_{a_1^B} p_{a_2^B}$ | $q_{a_1^A} p_{a_2^A} q_{a_1^B}$ | $q_{a_1^A} p_{a_2^A} p_{a_2^B}$ | $q_{a_1^A} p_{a_2^A}$ |
| 11 00 | $q_{a_1^A} q_{a_2^A} p_{a_1^B} p_{a_2^B}$ | 0 | 0 | 0 |
| 11 01 | $q_{a_1^A} q_{a_2^A} p_{a_1^B} q_{a_2^B}$ | $q_{a_1^A} q_{a_2^A} p_{a_1^B}$ | 0 | 0 |
| 11 10 | $q_{a_1^A} q_{a_2^A} q_{a_1^B} p_{a_2^B}$ | 0 | $q_{a_1^A} q_{a_2^A} p_{a_2^B}$ | 0 |
| 11 11 | $q_{a_1^A} q_{a_2^A} q_{a_1^B} q_{a_2^B}$ | $q_{a_1^A} q_{a_2^A} q_{a_1^B}$ | $q_{a_1^A} q_{a_2^A} q_{a_2^B}$ | $q_{a_1^A} q_{a_2^A}$ |

**Table S2:** Explanation of how to calculate 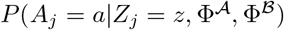 for all possible combinations of *z*_1_, *z*_2_ and *a*. It is assumed that alleles from different populations can not be IBD.

| $a$ | $z_1 = 0 \ z_2 = 0$ | $z_1 = 1 \ z_2 = 0$ | $z_1 = 0 \ z_2 = 1$ | $z_1 = 1 \ z_2 = 1$ |
| --- | --- | --- | --- | --- |
| $a_1^A \neq a_1^B \ a_2^A \neq a_2^B$ | $\Phi_{a_1^A, a_2^A}^A \Phi_{a_1^B, a_2^B}^B$ | 0 | 0 | 0 |
| $a_1^A = a_1^B \ a_2^A \neq a_2^B$ | $\Phi_{a_1^A, a_2^A}^A \Phi_{a_1^B, a_2^B}^B$ | $\frac{\Phi_{a_1^A, a_2^A}^A \Phi_{a_1^B, a_2^B}^B}{\sum_{x=1}^K \sum_{y=1}^K \sum_{k=1}^K \Phi_{k,x}^A \Phi_{k,y}^B}$ | 0 | 0 |
| $a_1^A \neq a_1^B \ a_2^A = a_2^B$ | $\Phi_{a_1^A, a_2^A}^A \Phi_{a_1^B, a_2^B}^B$ | 0 | $\frac{\Phi_{a_1^A, a_2^A}^A \Phi_{a_1^B, a_2^B}^B}{\sum_{x=1}^K \sum_{y=1}^K \sum_{k=1}^K \Phi_{x,k}^A \Phi_{y,k}^B}$ | 0 |
| $a_1^A = a_1^B \ a_2^A = a_2^B$ | $\Phi_{a_1^A, a_2^A}^A \Phi_{a_1^B, a_2^B}^B$ | $\frac{\Phi_{a_1^A, a_2^A}^A \Phi_{a_1^B, a_2^B}^B}{\sum_{x=1}^K \sum_{y=1}^K \sum_{k=1}^K \Phi_{k,x}^A \Phi_{k,y}^B}$ | $\frac{\Phi_{a_1^A, a_2^A}^A \Phi_{a_1^B, a_2^B}^B}{\sum_{x=1}^K \sum_{y=1}^K \sum_{k=1}^K \Phi_{x,k}^A \Phi_{y,k}^B}$ | $\frac{\Phi_{a_1^A, a_2^A}^A \Phi_{a_1^B, a_2^B}^B}{\sum_{k_1=1}^K \sum_{k_2=1}^K \Phi_{k_1, k_2}^A \Phi_{k_1, k_2}^B}$ |

## References

[1] Shaun Purcell et al. “Plink: A Tool Set for Whole-Genome Association and Population-Based Linkage Analyses”. In: American journal of human genetics 81 (Oct. 2007), pp. 559–75. doi: 10.1086/519795.

[2] Kermit Ritland. “Estimators for pairwise relatedness and individual inbreeding coefficients”. In: Genetical Research 67.2 (1996), pp. 175–185. doi: 10.1017/S0016672300033620.

[3] Brook G. Milligan. “Maximum-Likelihood Estimation of Relatedness”. In: Genetics 163.3 (2003), pp. 1153–1167. issn: 0016-6731. eprint: https://www.genetics.org/content/163/3/1153.full.pdf. url: https://www.genetics.org/content/163/3/1153.

[4] EA Thompson. “The estimation of pairwise relationships”. In: Ann. Hum. Genet. 39 (1975), pp. 173–188. doi: https://doi.org/10.1111/j.1469-1809.1975.tb00120.x.

[5] Anders Albrechtsen et al. “Relatedness mapping and tracts of relatedness for genome-wide data in the presence of linkage disequilibrium”. In: Genetic Epidemiology 33 (2008), pp. 266–274. doi:https://doi.org/10.1002/gepi.20378.

[6] Ida Moltke and Anders Albrechtsen. “RelateAdmix: a software tool for estimating relatedness between admixed individuals”. In: Bioinformatics 30.7 (Nov. 2013), pp. 1027–1028. issn: 1367-4803. doi: 10.1093/bioinformatics/btt652. eprint: https://academic.oup.com/bioinformatics/articlepdf/30/7/1027/17147070/btt652.pdf. url: https://doi.org/10.1093/bioinformatics/btt652.

[7] RV Rohlfs, SM Fullerton, and BS Weir. “Familial identification: population structure and relationship distinguishability”. In: PLoS Genetics 8 (2012). doi: 10.1371/journal.pgen.1002469.

[8] T. Thornton et al. “Estimating Kinship in Admixed Populations”. In: The American Journal of Human Genetics 91 (2012), pp. 122–138. doi: 10.1016/j.ajhg.2012.05.024.

[9] Ani Manichaikul et al. “Robust relationship inference in genome-wide association studies”. In: Bioinformatics 26.22 (Oct. 2010), pp. 2867–2873. issn: 1367-4803. doi: 10.1093/bioinformatics/btq559. eprint: https://academic.oup.com/bioinformatics/article-pdf/26/22/2867/16896963/btq559.pdf. url: https://doi.org/10.1093/bioinformatics/btq559.

[10] Rasmus Nielsen et al. “SNP calling, genotype calling, and sample allele frequency estimation from new-generation sequencing data”. English. In: PLOS ONE 7.7 (2012). e37558. issn: 1932-6203. doi: 10.1371/journal.pone.0037558.

[11] Thorfinn Sand Korneliussen and Ida Moltke. “NgsRelate: a software tool for estimating pairwise relatedness from next-generation sequencing data”. In: Bioinformatics 31.24 (Dec. 2015), pp. 4009–4011. issn: 1367-4803, 1367-4811. doi: 10.1093/bioinformatics/btv509. url: http://dx.doi.org/10.1093/bioinformatics/btv509.

[12] Kristian Hanghøj et al. “Fast and accurate relatedness estimation from high-throughput sequencing data in the presence of inbreeding”. en. In: GigaScience 8.5 (May 2019). issn: 2047-217X. doi: 10.1093/gigascience/giz034.

[13] Heng Li et al. “The Sequence Alignment/Map format and SAMtools”. In: Bioinformatics (Oxford, England) 25.16 (2009), pp. 2078–2079. issn: 1367-4803. doi: 10.1093/bioinformatics/btp352. url: https://europepmc.org/articles/PMC2723002.

[14] Thorfinn Sand Korneliussen, Anders Albrechtsen, and Rasmus Nielsen. “ANGSD: Analysis of Next Generation Sequencing Data”. In: BMC bioinformatics 15 (Nov. 2014), p. 356. issn: 1471-2105. doi: 10.1186/s12859-014-0356-4. url: http://dx.doi.org/10.1186/s12859-014-0356-4.

[15] L. Skotte, T. S. Korneliussen, and A. Albrechtsen. “Estimating Individual Admixture Proportions from Next Generation Sequencing Data”. In: Genetics 195.3 (2013), pp. 693–702. doi:https://doi.org/10.1534/genetics.113.154138.

[16] Jonas Meisner and Anders Albrechtsen. “Inferring Population Structure and Admixture Proportions in Low-Depth NGS Data”. English. In: Genetics 210.2 (2018), pp. 719–731. issn: 0016-6731. doi: 10.1534/genetics.118.301336.

[17] Ravi Varadhan and Christophe Roland. “Simple and Globally Convergent Methods for Accelerating the Convergence of Any EM Algorithm”. In: Scandinavian Journal of Statistics 35.2 (June 2008), pp. 335–353. issn: 0303-6898, 1467-9469. doi: 10.1111/j.1467-9469.2007.00585.x. url: http://doi.wiley.com/10.1111/j.1467-9469.2007.00585.x.

[18] 1000 Genomes Project Consortium et al. “A global reference for human genetic variation”. en. In: Nature 526.7571 (Oct. 2015), pp. 68–74. issn: 0028-0836, 1476-4687. doi: 10.1038/nature15393. url: http://dx.doi.org/10.1038/nature15393.

[19] D.H. Alexander, J. Novembre, and K. Lange. “Fast model-based estimation of ancestry in unrelated individuals”. In: Genome Research 19 (2009), pp. 1655–1664. doi: 10.1101/gr.094052.109.

[20] A McKenna et al. “The Genome Analysis Toolkit: a MapReduce framework for analyzing next-generation DNA sequencing data”. In: Genome Res 20.9 (Sept. 2010), pp. 1297–1303. doi: 10.1101/gr.107524.110. url: https://www.ncbi.nlm.nih.gov/pubmed/20644199?dopt=Abstract.

[21] Shaun Purcell et al. “PLINK: a tool set for whole-genome association and population-based linkage analyses”. en. In: American journal of human genetics 81.3 (Sept. 2007), pp. 559–575. issn: 0002-9297. doi: 10.1086/519795. url: http://dx.doi.org/10.1086/519795.

[22] T. J. Pemberton et al. “Inference of Unexpected Genetic Relatedness among Individuals in HapMap Phase III”. In: American journal of human genetics 87.4 (2010), pp. 457–464. doi:https://doi.org/10.1016/j.ajhg.2010.08.014.

[23] Elizabeth A Thompson. “Identity by descent: variation in meiosis, across genomes, and in populations”. en. In: Genetics 194.2 (June 2013), pp. 301–326. issn: 0016-6731, 1943-2631. doi: 10.1534/genetics.112.148825. url: http://dx.doi.org/10.1534/genetics.112.148825.

[24] Carl Veller et al. “Variation in genetic relatedness is determined by the aggregate recombination process”. en. May 2020. url: https://www.biorxiv.org/content/10.1101/2020.05.25.115048v1.

[25] Emil Jørsboe, Kristian Ebbesen Hanghøj, and Anders Albrechtsen. “fastNGSadmix: admixture proportions and principal component analysis of a single NGS sample”. English. In: Bioinformatics 33.19 (Oct. 2017), pp. 3148–3150. issn: 1367-4803. doi: 10.1093/bioinformatics/btx474.

[26] B G Lindsay. “Composite likelihood methods”. In: Contemporary Mathematics (1988). issn: 0271-4132. url: https://ci.nii.ac.jp/naid/20000924949/.

